# Solvent-specific bioactivities of cone, leaf, and stem extracts from a native Finnish wild hop

**DOI:** 10.64898/2026.03.26.714411

**Authors:** Lidija Bitz, Juha-Matti Pihlava, Pertti Marnila, Lucia Blasco, Ville Paavilainen, Merja Hartikainen, Anna Nukari, Dale Tranter, Teija Tenhola-Roininen

## Abstract

The genetically authenticated Finnish hop genotype LUKE 2541 obtained from wild was evaluated for antibacterial, anti-inflammatory, and anticancer activities. Water extracts from hop cones inhibited the Gram-positive bacteria *Staphylococcus aureus* and *Bacillus cereus*, with MIC values of 0.094– 0.188⍰mg/mL, whereas Gram-negative strains showed limited sensitivity. In LPS-primed THP-1 cells, both IPA and IPA-Control extracts reduced reactive oxygen species formation in a dose-dependent manner, exhibiting similar IC_50_ values (50.41⍰µg/mL and 35.41⍰µg/mL). This hop genotype also displayed clear tissue- and solvent-dependent antiproliferative effects in human cancer cell lines. Bioactivity was strongly enriched in hop cones and predominantly associated with non-polar extracts, particularly hexane and dichloromethane fractions, which produced marked, dose-dependent reductions in cell viability. In contrast, aqueous and methanolic extracts were largely inactive, underscoring the critical importance of extraction chemistry and tissue selection. Sensitivity varied among cancer cell lines, with colorectal cells generally more responsive and leukemia cells less affected, highlighting cell-specific susceptibility. Further research is needed to elucidate underlying mechanisms, determine selectivity toward non-malignant cells, and identify the active compounds responsible for all in all investigated effects.

## INTRODUCTION

Hop (*Humulus lupulus* L.) is climbing dioecious plant belonging to the Cannabaceae family and thus being related to the cannabis and nettle. The cone-like female inflorescences (hop cones) have been recognized and exploited in traditional practices and natural folk medicine since ancient times (Karabin et al., 2015). Traditional knowledge about the use of hops is abundant and diverse, blood purification, combating anxiety and insomnia, leprosy as well as fabric and paper production being only some of the pre-historical applications. Defining hop as a medicinal plant might come as a surprise to many as during the last centuries this plant has been vastly exploited and best known only in the beer and brewing industry. Resins from hop cones lupulin glands are essential ingredients to beer. Synthesized alpha acids (mainly humulon and lupulon) give beer its bitterness, while strong smelling terpens give beers their unique aroma. Not only does hop confer bitterness and aroma, but also it acts as a natural conservative via phenylpropanoid derivatives, which exhibit toxicity against microbes. However as already known from distant past, besides the exploitation of hop cones for the brewing industry, hop offers alternative and supplemental uses because it is also extraordinarily rich in flavonoids (the major one being xanthohumol), which is highly valued for being an active anti-tumor agent. Conversion of flavonoids facilitates beer enrichment in phytoestrogens (e.g., 8-prenylnaringenin has been characterized as one of the most potent phytoestrogens isolated until now) (Karabin et al., 2015). Considering bioactive properties of certain hop compounds, this plant could be regarded as a strong candidate for molecular farming exploitation («biopharming») and through *in vitro* culture. Biosynthetic pathways of hop metabolites related to human benefits are well documented. Research is also orientated toward elucidation of the metabolism of flavonoids and establishing biological activity of their metabolites. These are seen as in its anti-infective properties, anti-cancer activities, and possible preventative activity towards Alzheimer disease and many more. Up to now there has been only rising evidence of benefits to wide range of diverse aspects of human health and well-being. Foodborne illnesses occur because of consumption of contaminated foods or beverages, and microbial pathogens are known to be the most probable cause, bacteria being the first cause, and viruses the second (WHO, 2015). The use of natural bio-preservation methods could ensure the safety and food quality while avoiding chemicals, synthetic compounds or other thermal or radiation methods. Consumers are more aware of health implications of undesirable chemicals, thus a trend to evaluate natural antimicrobials is quickly arising (Gyawali and Ibrahim, 2014; Quinto et al., 2019). Also concerns like the increase of multidrug-resistant microorganisms would support the active search for natural antimicrobials.

A phylogenetic analysis indicated that hop was introduced to Europe from its putative place of origin in China about one million years ago (Murakami et al., 2006). In Finland there is historical bases (hop is a native plant to southern and central Finnish shores and broad-leaved forests on stream banks. Hop began to grow inland at a time when Finland’s coastline was washed by the Litorina Sea, some 7,000– 8,000 years ago) and a tradition of hop production (its inflorescence was a means of paying tax when Finland was part of Sweden, and peasants were legally obliged to cultivate hops until 1915. There was a time when every croft and small cottage had hop trellis in the yard. Later there were even hundreds of commercial hop fields for the brewery industry – the Finnish equivalent of southern European vineyards). Hop production is nowadays represented almost only by individual noncommercial work done by enthusiastic people.

Pollen evidence indicates hop has been present since at least 8500 BC, long before cultivation began ca. 5300–4000 BC (Alenius et al., 2013). Historically, hop was used not only for brewing but also as medicine, food, fibre, and household materials. Cultivation declined in the 19th century with the rise of imported cultivars, but interest in local hops has resurged alongside the growth of microbreweries and recognition of bioactive compounds such as phytoestrogens and flavonoids. This has prompted efforts to identify and select native genotypes for renewed cultivation.

Broad set of samples representing Finnish hops have been genetically and chemically evaluated previously. The goal of this study is to deepen the understanding of Finnish hops compositions that goes beyond use of hops in flavoring the beers. In the present study, we evaluated true-to-type Finnish hop regarding their chemical composition as well their antimicrobial, anti-inflammatory and anticancer activities. We discuss the evaluated properties and propose future research in this field based on the results obtained of this study and comparative analyses with previous related research.

## MATERIALS AND METHODS

### Selection and sampling of hop plant materials

Based on our previous research (Bitz et al., 2021) we have selected representative true to type wild hop natively growing in Finland for thousands of years. One hop genotype (LUKE-2541) has been selected, and leaves, cones and stems have been sampled for extract preparation. HOP LUKE-2541 has been genetically (microsatellite fingerprinting and genetic analyses) and chemically (α-acids - cohumulone, n-humulone and adhumulone and β-acids - colupulone, n-lupulone + adlupulone (AdL) previously characterized as described in (Bitz et al., 2021; Luong et al., 2025).

### Extract preparations

Leaf, stem and cone samples have been obtained from mature hop plant (LUKE-2541) at cone maturity stage and extracts prepared as described below:

Control lager: 1,5 g tissues homogenized in 1 l MilliQ water by Bamix. Stirred in magnetic stirrer overnight at room temperature. Next morning sifting through tea sift and Retsch sifter (450 um, 40 mesh). Freezing part of the liquid and freeze-drying the rest.

Lager boil: 1,5 g tissues homogenized in almost boiling 1 l MilliQ water by hand blender. Mild boiling for 1 h. Heat off. Additional homogenization by hand blender. Post-boil stand on hot plate for 0,5 h. Cooling 1,5 h in 1-5 °C to cool down to room temp, storage overnight at room temperature without mixing. Next morning sifting through tea sift and Retsch sifter (450 um, 40 mesh). Freezing part of the liquid and freeze-drying the rest.

Control IPA: 6 g tissues homogenized in 1 l MilliQ water by Bamix. Stirred in magnetic stirrer overnight at room temperature. Next morning sifting through tea sift and Retsch sifter (450 um, 40 mesh). Freezing part of the liquid and freeze-drying the rest.

IPA boil: 3 g tissues homogenized in almost boiling 1 l MilliQ water by hand blender. Mild boiling for 1 h. Heat off. Addition of 3 g hops and homogenization by hand blender. Post-boil stand on hot plate for 0,5 h. Cooling 1,5 h in 1-5 °C to cool down to room temp, storage overnight at room temperature without mixing. Next morning sifting through tea sift and Retsch sifter (450 um, 40 mesh). Freezing part of the liquid and freeze-drying the rest.

### Anti-bacterial analyses

The experiment was carried out on two species of gram-positive bacteria *Staphylococcus aureus* ATCC 33862 and *Bacillus cereus* ATCC 14579 and two-gram negative species, *Escherichia coli* ACTT25922 and *Pseudomonas aeruginosa* ACTT 27853. The hop extracts were diluted in water using two-fold methods and examined at concentrations ranging from 6 to 0.188mg/mL for different extraction methods. The minimal inhibitory concentration (MIC) of bacterial growth was determined by inoculating the bacterial culture with the broth Mueller Hinton (MH) or Luria Broth (LB) at different hop concentrations. The turbidity of each bacterial suspension was adjusted to obtain 1.10^4^ colony-forming units (CFU)/mL in the growth media, before inoculation. Hop extracts were filtered through a 0,22µm pore filter and dispensed into wells using sterile pipette tips, and sterile water was added to reach different concentrations and dispensed in triplicate into the microtiter wells. The inhibition was determined after incubation at 37°C for 24 h. A positive growth control without hop extract added was included (only optimal growth medium). The lowest hop concentration required to inhibit bacterial growth was defined as the MIC. All MIC measurements were carried out in duplicate.

### Anti-inflammatory evaluations

#### Reagents

Stock solution of 20 mM luminol (5-amino-2,3-dihydro-1,4-pthalazinedione; ICN Biomedicals, Aurora, OH, USA) was made in 0.2 M sodium borate buffer (pH 9.0). Solution was warmed up to 50°C and shaken until the crystals were dissolved. Zymosan powder (zymosan A from yeast *Saccharomyces cerevisiae*; Sigma-Aldrich) was autoclaved (120°C for 20min) in phosphate buffered saline (PBS, pH 7.4) in concentration of 10 mg/mL, washed three times with centrifugation (10 min at 1000g) in HBSS buffer (Hank’s balanced salt solution, without phenol red; pH 7.4) and finally an 10 mg/mL solution of zymosan was made in HBSS buffer. The zymosan was opsonized by pooling fresh human sera from three healthy adult volunteers and by incubating the zymosan with serum pool in water bath for 30 min at 37°C with continuous gentle mixing. The opsonization mixture consisted of 50% serum and of 50% (v/v) zymosan solution (10 mg/ml). The opsonized zymosan (OZ) was then washed three times with HBSS buffer as described above and suspended in HBSS (5 mg/mL) and divided into 1-2mL aliquots and stored at -80°C. Lipopolysaccharide (LPS) from *Escherichia coli* 055:B5 (LPS; Sigma-Aldrich) and horseradish peroxidase (HRP; Merck 10814407001 Roche, Germany) were dissolved in HBSS buffer to obtain 2.5 mg/ml and 800 U/mL solutions, respectively. LPS stock solution was divided into aliquots and stored at -22°C. HRP solution HRP was prepared weekly by dissolving into PBS (pH 7.4) and was kept at +4°C maximum five days. 10x Resazurin stock solution (44µM) was prepared by dissolving resazurin sodium salt (R7017, Sigma-Aldrich) in PBS and mixing vigorously, followed by filter sterilization through 0.2µm filter. Resazurin stocks are stored at 4°C and protected from light exposure by aluminum foil.

#### Sample preparation

For immunological tests the pH’s of the hop tea samples IPA and IPA Control were adjusted to physiological pH 7.4 by adding carefully to 8mL of HOP tea samples 0.1N NaOH and HBSS buffer to reach 10mL volume (80% of tea and 20% of buffers, v/v). The samples were then centrifuged by 4500 ***g*** for 15min to remove undissolved material and finally put through 0.20µm filter to remove microparticles that might activate leukocytes. The dry matters were measured by weighting after drying at 98 °C for 1.5 hours. Before removing the undissolved material, the dry matter contents of IPA and IPA Control were 1.82mg/mL and 1.42mg/mL, respectively, and after centrifugation and filtering the dry matters of IPA and IPA Control were 1.062mg/mL and 0.800 mg/mL. These dry matter values do not contain the salt added in pH adjustment. The proportions of undissolved material were 27.1% in IPA and 29.6% (w/w). The pH adjusted and centrifuged HOPS tea filtrates were used in leukocyte tests and stored in aliquots at -22°C. The final test concentrations ranged from 0.42 to 303 µg/mL (w/v) in IPA and from 0.31 to 229 µg/mL (w/v) in IPA Control.

#### Respiratory burst measurements

Kinetic measurements of THP-1 cell respiratory burst (RB) activity were conducted as described in Tompa et al. (2011) with some modifications. For measurements microplate reader (Hidex Sense, idex LTD.Turku, Finland) and white 96-well flat bottom non treated microtiter plates (Nunc 36105, Roskilde, Denmark) and THP-1 cells in passages 5–30 after thawing were used. The cells were washed once by centrifuging at 300***g*** for 5 min and gently suspended into gel-HBSS (HBSS buffer supplemented with 0.1% w/v gelatin, gelatin A from swine skin, Sigma-Aldrich) at room temperature (RT) the concentration varying between 5 to 9 × 10^5^ cells/ml.

The THP-1 cells (around 3-6 × 10^5^ /ml) were incubated in microtiter plate at 37°C for 15 min in temperature-controlled chamber with different concentrations of extracts. Then LPS solution was added to prime the cells into inflammatory state, and the incubation was continued for 35 min at 37°C the LPS concentration being 10µg/ml. After altogether 50 min preincubation with extracts the plate was quickly put into luminometer and kept in luminometer measuring chamber with oversteering temperature control for 5 min to stabilize the sample temperature to 37°C. The reaction was started by promptly adding opsonized zymosan together with luminol and HRP in 50µl volume. During RB reaction the final concentrations of HRP was 25 U/ml, luminol 0.5 mM and opsonized zymosan 250 mg/ml. The chemiluminescence (CL) signal was counted and integrated for 0.5 s at 2.33 min intervals for 90 min to obtain kinetic curves. The CL peak (relative light units, rlu) was regarded as the CL value of the sample. The efficacies of extracts were expressed as IC50-values i.e. concentrates which inhibit 50% of the CL emission obtained from cells without extracts on same plate. In dose response curves the concentrations of extracts are those during the incubation with cells and LPS. The experiments were repeated 8 times, and one IPA and IPA-Control was used.

Leukemic promonocytes have different properties from native peripheral blood monocytes. Since THP-1 cells do not express myeloperoxidase, HRP adding was needed as a substitute to obtain a CL emission with sufficient intensity. Luminol CL emission from leukocytes is solely dependent on peroxidase activity.

#### Cell culture

THP-1 human promonocytes leukemic cell line (Leibniz Institute, DMSZ German Collection of Microorganisms and Cell Cultures) was cultured in RPMI 1640-GlutaMAX^™^ medium (Gibco^®^ Invitrogen^™^, USA) supplemented with 10% of heat-inactivated (30min at 56°C) fetal bovine serum (FBS, Gibco), 10 mM HEPES (BioWhittaker, Lonza), 1 mM sodium pyruvate (Gibco), 2.0 g/L [+]D-glucose (Gibco), 0.5 mM 2-mercaptoethanol (Gibco), 2mM L-glutamine (Gibco), 50 U/ml penicillin and 50 µg /ml of streptomycin (Gibco). Cells were maintained in a humidified atmosphere with 5% CO_2_ at 37°C. Medium was changed every 3-4 days and cells were sub-cultured with resuspension of 2 × 10^5^ viable cells/mL and the density of viable cells was not allowed to rise higher than 9 × 10^5^ /ml. The viable cell concentration was counted by staining with 0.4% trypan blue solution and counted by microscope and Bürker chamber or with Countess II FL cell counter (Invitrogen).

### Anti-cancer evaluations

#### Inhibition of cancer cell line proliferation

HCT116 human colorectal carcinoma cells were cultured in McCoy’s 5A medium (Gibco^®^) supplemented with 10% fetal bovine serum (Gibco^®^). K562 human myeloid leukemia and NCI-H23 human lung adenocarcinoma cells were cultured in RPMI 1640 medium (prepared from ready mix, 1060122, ICN) supplemented with 10% fetal bovine serum (Gibco^®^). A549 human lung carcinoma cells were cultured in DMEM (prepared from ready mix, D7777, Sigma-Aldrich) supplemented with 10% fetal bovine serum (Gibco^®^). HCT116, NCI-H23 and A549 cells were passaged every 2-3 days to maintain cell confluence below 80%. K562 cells were passaged every 2-3 days to maintain cell density below 2 × 106 viable cells per ml. HCT116, NCI-H23, A549 and K562 cell concentrations were counted using a BioRad TC10 cell counter following staining with trypan blue solution.

#### Mammalian cell proliferation assays

Viable cell densities were estimated using an automated cell counter (BioRad TC10) before each experiment. 100µl of cells at a density of 25,000 cells/ml were seeded into black walled view plates (Viewplate-96, Perkin Elmer) and incubated for 24h at 37 °C and 5% CO_2_. 5x working solutions were prepared immediately prior to use by dilution of plant extracts into appropriate culture media, followed by serial dilutions into media containing an equivalent concentration of the respective carrier solvent. Cells were dosed by addition of 25µl of 5x working stocks and incubated at 37 °C and 5% CO_2_ for 72h.

To assess cell viability, 12.5µl of 10x resazurin stock was added per well, followed by a further 4h incubation at 37 °C and 5% CO_2_. Plates were then scanned by a microplate reader (Perkin Elmer Enspire) for 620 nm emission while subjected to 580 nm excitation. Curves were fitted to the data using GraphPad Prism 9, and calculated IC50 values plotted to a heat map.

## RESULTS

### Anti-bacterial

After an incubation period of 24 hours, bacterial growth was observed. Four bacterial strains were investigated for their resistance to water extracts obtained from hop cones. Antimicrobial activity was confirmed against the Gram-positive *S. aureus* and *B. cereus* for all hop cones extracts, with the mean MICs in the range of 0.094 mg/mL to 0.188 mg/mL.

### Anti-inflammatory

Both IPA and IPA-Control samples inhibited the reactive oxygen species (ROS) generation during the THP-1 cell respiratory burst in dose dependent manner (see Figure 1). The IC50 concentrations (inhibited 50% of ROS production) were 50.4 S.D. ±22.6 µg/mL in IPA and 35.4 S.D. ±29.8 µg/mL in IPA-Control. The IC50 values were not statistically different (P=0.86). Both samples inhibited most of the ROS in highest concentrations of 230 and 303µg/mL (see Figure 1).

**Figure 1:**
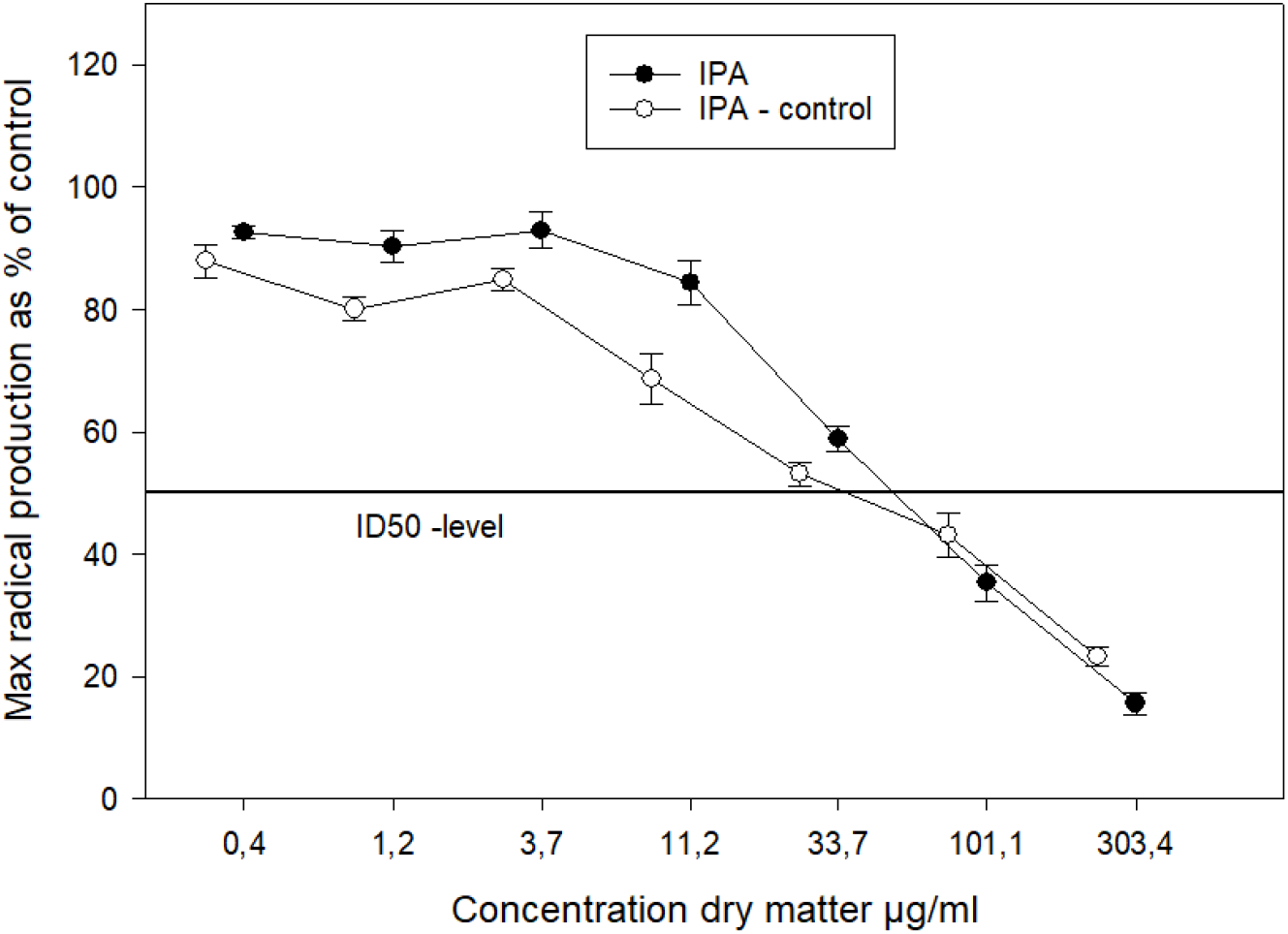
The effects of HOP tea extracts IPA and IPA-Control on maximum radical production rate of THP-1 cells during respiratory burst. The cells were primed to inflammatory state with *E. coli* LPS and 35 min later the respiratory burst reaction was induced by human serum treated yeast cell wall particles. The values are the peroxidase dependent luminol enhanced chemiluminescence reaction maxima presented as per cent of that of the controls with no extract. The error bars represent standard error of the mean (S.E., n=9). The ID50 is the level of 50% inhibition.

**Figure 2.**
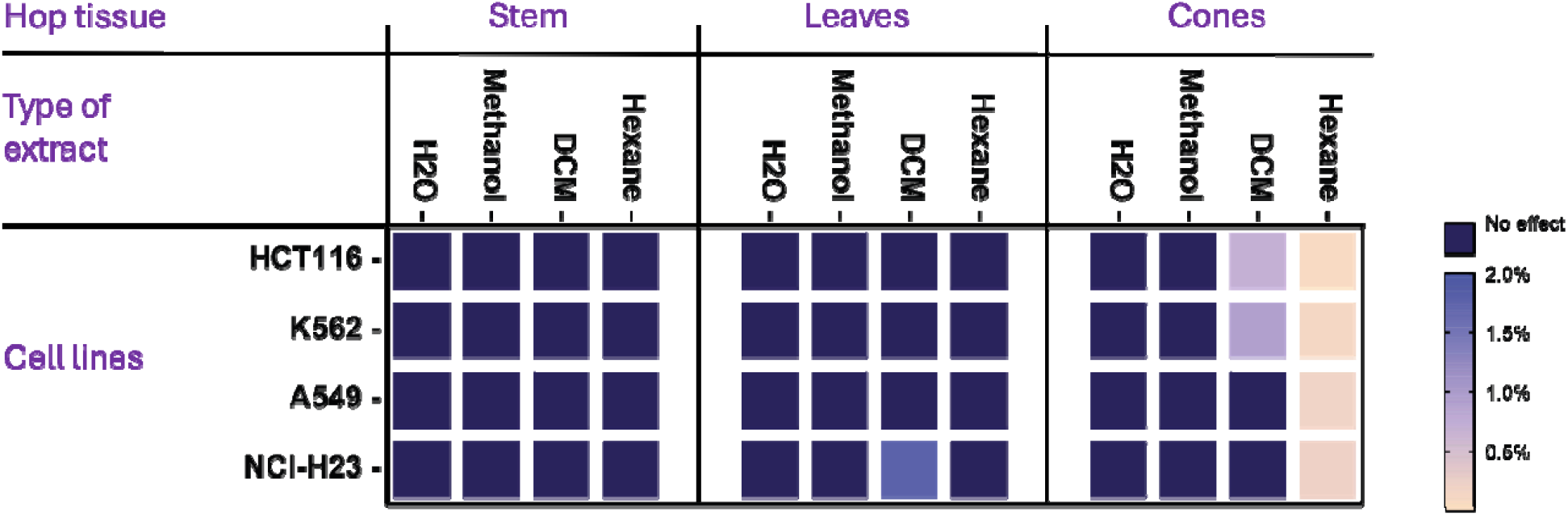
Heat map of tissue- and solvent-dependent antiproliferative activity of hop extracts in human cancer cell lines.

**Figure 3.**
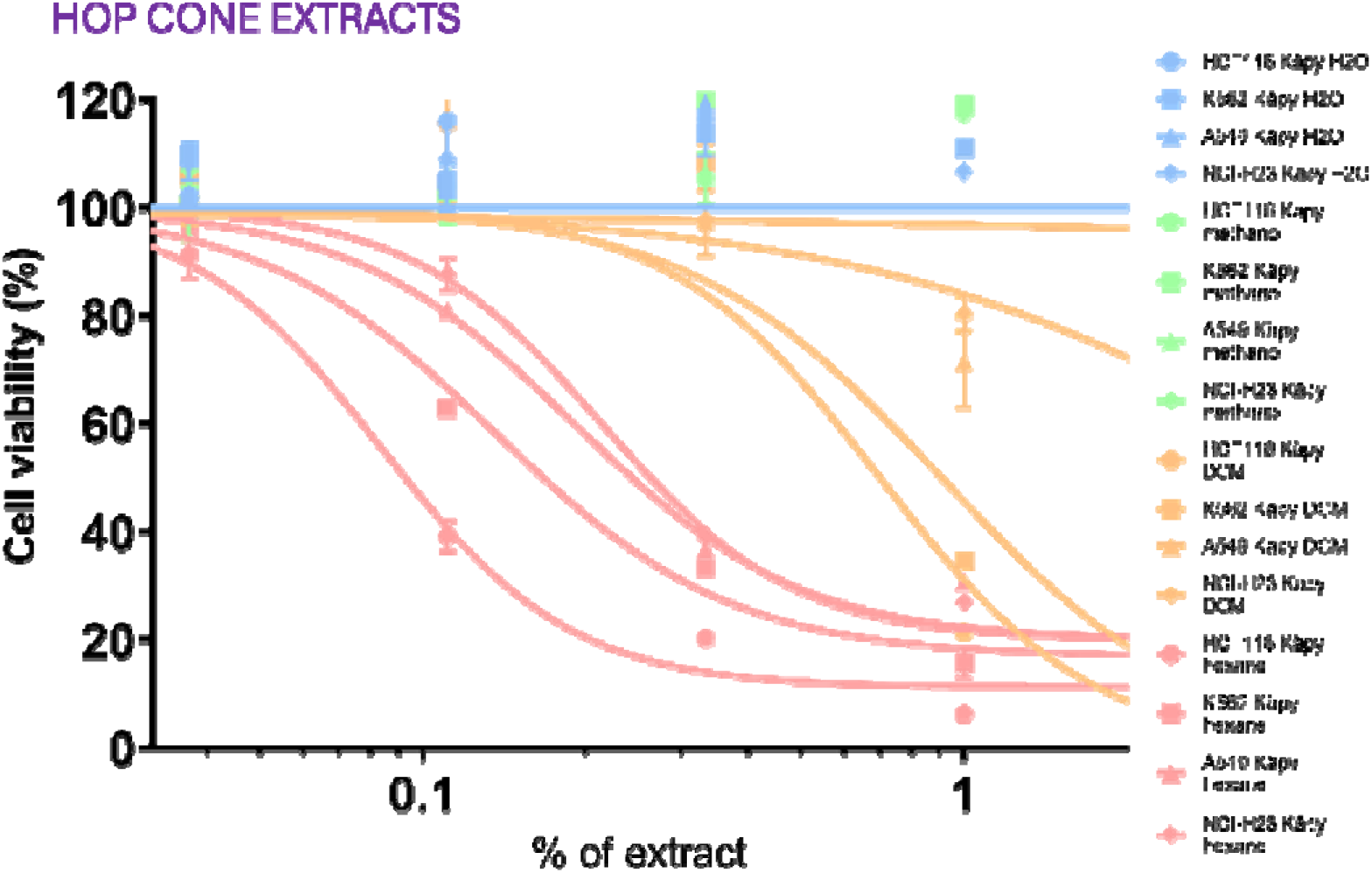
Comparative dose–response analysis of aqueous, methanolic, DCM, and hexane hop cone extracts on human cancer colorectal (HCT116), lung (A549 and NCI-H23) and leukemia (K562) cell lines.

### Inhibition of cancer cell line proliferation

Selected cancer cell lines with known unique molecular profiles were tested. A range of antiproliferative activities were observed for hop cone extracts prepared with either hexane or DCM. It appears there is a rank order of relative potencies between the different cell lines.

## DISCUSSION

### Anti-bacterial

Due to the global increase in multidrug resistance, the high of developing new antibiotics, and the time-consuming nature of the process, natural extracts have been proposed as alternative antimicrobial agents or as supportive treatments to traditional therapies. Previous studies have shown significant antibacterial activity of hop extract against different Gram-positive pathogenic bacteria (Di Lodovico et al., 2020).

One of the most common foodborne bacteria causing food poisoning is *Staphylococcus aureus*. These Gram-positive bacteria are commonly found on the skin, throats and nostrils of healthy people and animals, and it does not cause illness unless it is transmitted to food products where it can grow and produce toxins, but they can be also methicillin-resistant (MRSA) and vancomycin-resistant (VRSA) which emphasizes the need for continue research on new antibiotics.

The use of hops in beer manufacturing serves as a natural way of preserving the quality of the liquid because of the bitter acids that inhibit especially Gram-positive bacterial growth. Despite the extensive knowledge of antibacterial activities of hops (Kramer et al., 2015); more and more, there is an increasing need to continue screening for antimicrobial potential of individual hop compounds as well as different extraction techniques.

**Table 1.** Minimal inhibitory concentrations (MICs) of water extracts from the hop cones collected in 2019 in Finland against bacteria.

| Hop | Extraction method | MIC (mg/L hop extract from cones) |  |  |  |
| --- | --- | --- | --- | --- | --- |
|  |  | *B. cereus* | *S. aureus* | *E. coli* | *P.aeruginosa* |
| 19-862 | 001 | 0,750 | 0,750 | 1,500 | 1,500 |
|  | 002 | 0,750 | 0,188 | 1,500 | 1,500 |
|  | 003 | 3,000 | 3,000 | 6,000 | 6,000 |
|  | 004 | 0,188 | 0,188 | 3,000 | 3,000 |
| 19-863 | 001 | 0,094 | 0,750 | 1,500 | 1,500 |
|  | 002 | 0,094 | 0,094 | 1,500 | 1,500 |
|  | 003 | 0,375 | 3,000 | 6,000 | 6,000 |
|  | 004 | 0,188 | 0,188 | 3,000 | 3,000 |

### Anti-inflammatory

THP-1 promonocytes are widely and extensively used models for human monocytes to study monocyte/macrophage functions, mechanisms, signaling pathways and nutrient transport. THP-1 cells resemble normal monocytes with respect to many criteria such as morphology, secretory products, oncogene expression, membrane antigens and expression of genes involved in lipid metabolism (Auwerx, 1991; Chanput et al., 2014). We used a LPS primed THP-1 cell model (Tompa et al., 2011) to study the effects of the HOPS extracts produced on the oxygen radical production of activated human THP-1 promonocytes during respiratory burst process.

Human serum opsonized zymosan particles used for triggering respiratory burst are recognized in THP-1 cells by the CD11b/CD18 (CR3) receptor complex together with Toll-like receptors 2 and possibly 4. THP-1 cells constitutively express serum opsonin recognizing receptors including complement receptor 1 (CD35) And immunoglobulin recognizing Fcg-receptors 1 (CD64) and 2 (CD32), although in lesser amounts than in native peripheral blood monocytes. These receptors provide an essential link between humoral and cellular immune systems by functioning as key molecules for phagocytosis, adhesion and triggering the release of inflammatory mediators e.g. IL-1β and TNF-α. Chronic overactivity of this system contributes to development of inflammation related widespread diseases. Thus, free radicals from activated phagocytes are of utmost importance due to their role in the development of numerous pathological states including atherosclerosis, age-related neurological disorders and type II diabetes. Strategies that target cell priming and overactivity would not only be expected to prevent disease initiation but may also have a therapeutic role, as monocyte–macrophage recruitment is a dynamic process that occurs throughout disease progression. Food compounds having anti-inflammatory properties may have beneficial effects about cardiovascular disease and type II diabetes (Jia et al. 2014, Castillo et al. 2020).

In this cell model the IPA and IPA-Control samples inhibited radical production by phagocytes. It suggests that components in HOP teas could possibly inhibit also *in vivo* harmful physiological processes mediated by long lasting inflammation. This result is in concordance with the recent studies by (Caban et al., 2020) where a polyphenol rich extract from HOPS suppressed expression of pro-inflammatory molecules IL-6, COX-2 mRNA and NF-κB in LPS activated murine macrophages (RAW 264.7). *In vivo* mouse model of hop derived iso-alpha acid preparation (freeze dried isohumulone potassium salt) added into Western type diet (500mg/kg body weight for 15 weeks) inhibited different pathophysiological steps in development of non-alcoholic fatty liver disease (Mahli et al., 2018). The results indicated reduced oxidative stress, pro-inflammatory gene expression and immune cell infiltration to hepatic tissue. Together with previous studies our results with THP-1 suggest that hop derived compounds have promising beneficial effects in prevention of Western diet related to widespread lifestyle diseases.

### Inhibition of cancer cell line proliferation

This study demonstrates that a genetically authenticated Finnish hop genotype (LUKE-2541) exhibits clear tissue- and solvent-dependent antiproliferative activity in human cancer cell lines. Bioactivity was strongly enriched in hop cones and confined mainly to non-polar extracts, particularly hexane and dichloromethane fractions, which induced pronounced, concentration-dependent reductions in cell viability. Aqueous and methanolic extracts were largely inactive, highlighting the importance of extraction chemistry. These findings are consistent with prior reports identifying hydrophobic hop constituents, such as bitter acids and prenylated flavonoids, as key mediators of anticancer activity through modulation of apoptosis and inflammatory signalling pathways (Girisa et al., 2021; Knez Hrnčič et al., 2019). Furthermore, the weak activity of brewing-style aqueous extracts aligns with evidence that thermal processing and polarity influence hop bioactivity (Busch et al., 2015). Together, these results support Finnish native hops as a promising bioactive resource beyond brewing.

Differences in sensitivity among cancer cell lines indicate cell-specific susceptibility to hop extracts, likely influenced by variations in metabolic capacity, membrane permeability, and intracellular signalling networks. Colorectal cancer cells tended to exhibit stronger responses, whereas leukemia cells were generally less sensitive, highlighting the importance of considering cancer type when evaluating plant-derived bioactives. While the present study focused on antiproliferative outcomes, further investigation is required to define underlying mechanisms, assess selectivity against non-malignant cells, and validate compound-specific effects.

## Conclusions

The results indicated reduced oxidative stress, pro-inflammatory gene expression and immune cell infiltration to hepatic tissue. Together with previous studies our results with THP-1 cells suggest that hop derived compounds have promising beneficial effects in the prevention of Western diet-related widespread lifestyle diseases.

The extract obtained from dried hop flowers showed good antimicrobial activity against the tested Gram-negative strains. As a result, those hops could be useful in the creation of an antibacterial product based on natural compounds with numerous industrial applications. However, more research is needed to describe the chemical composition and amounts of compounds in Finish wild hops that contribute to the antimicrobial and antioxidant activity.

Finnish hop genotype LUKE-2541 harbours potent, solvent-extractable antiproliferative activity localized to cones, with hexane⍰>⍰DCM ≫ methanol ≈ water as a consistent pattern across four human cancer cell lines. The present work positions Finnish native hops as a promising resource for bioactive discovery and provides a clear roadmap for compound-level identification and mechanistic validation.

## Acknowledgement

This study has been established through the Natural resources institute Finland strategic funding and financed by Finnish Ministry of Agriculture and Forestry (MMM), Maiju and Yrjö Rikkala Garden Foundation and PBL Brewing Laboratory.

